# The Anti-Epileptic Ketogenic Diet Alters Hippocampal Transporter Levels and Reduces Adiposity in Aged Rats

**DOI:** 10.1101/178319

**Authors:** Abbi R. Hernandez, Caesar M. Hernandez, Keila T. Campos, Leah M. Truckenbrod, Yasemin Sakarya, Joseph A. McQuail, Christy S. Carter, Jennifer L. Bizon, Andrew P. Maurer, Sara N. Burke

## Abstract

Nutritional ketosis is induced by high fat/low carbohydrate dietary regimens, which produce high levels of circulating ketone bodies, shifting metabolism away from glucose utilization. While ketogenic diets (KD) were initially introduced to suppress seizures, they are garnering attention for their potential to treat a myriad of neurodegenerative and metabolic disorders that are associated with advanced age. The feasibility and physiological impact of implementing a long-term KD in old animals, however, has not been systematically examined. In this study, young and aged rats consumed a calorically- and nutritionally-matched KD or control diet for 12 weeks. All KD-fed rats maintained higher levels of β-hydroxybutyrate (BHB) and lower levels of glucose relative to controls. However, it took the aged rats longer to reach asymptotic levels of BHB compared to young animals. Moreover, KD-fed rats had significantly less visceral white and brown adipose tissue than controls without a loss of lean mass. Interestingly, the KD lead to significant alterations in protein levels of hippocampal transporters for monocarboxylates, glucose and vesicular glutamate and GABA. Most notably, the age-related decline in vesicular glutamate transporter expression was reversed by the KD. These data demonstrate the feasibility and potential benefits of KDs for treating age-associated neural dysfunction.

## Introduction

The past decade has seen a surge in the number of studies investigating the therapeutic potential of ketogenic diets (KDs), which shift the body’s primary metabolic substrate from glucose to fat, to treat a wide range of neurological disorders. Clinically, KDs have been well documented to improve brain functioning in epilepsy(1,2), as well as reduce cognitive dysfunction in rodent models of seizure disorders(3). While the KD has been used as a treatment for epilepsy for nearly a century(4,5), more recently it has been implicated as an effective therapeutic in several other neurological disease states including Parkinson’s Disease, Alzheimer’s Disease and dementia(6,7), as well as diabetes, cancer, and cardiovascular disease(7–9). The biggest risk factor for many of the diseases for which the KD is currently being investigated is advancing age. However, a well-controlled, systematic study as to how age affects the ability to enter and maintain a state of ketosis, as well as if this causes any adverse effects over the long term, has yet to be documented.

Currently, most studies involving rodent models of age-related neurodegenerative diseases have tested the effects of nutritional ketosis in young animals(6,10,11). Because of the age-related changes in metabolic function and capacity, there is great potential for differential effects across the lifespan. Thus, there is a direct need for a KD study in an aged rodent model that examines the physiological and biochemical impact of long-term nutritional ketosis. Moreover, previous studies using KDs in rodents are hard to reconcile and often report conflicting results due to the high degree of inconsistency in dietary implementation across studies (See Table 1). Supplementation with medium chain triglycerides, which quickly enter circulation and are readily converted into ketones by the liver, have shown to have modest effects on hippocampal morphology(12) and behavior(13). While these data demonstrate the feasibility of KD-based therapies, prior MCT-based dietary regimes have not been calorically matched across ketogenic and control groups. Moreover, young ketogenic animals were not included in the experimental designs of previously reported studies, so it is not possible to determine if the reported effects are specific to aged animals or a ubiquitous impact of ketosis.

**Table 1:**
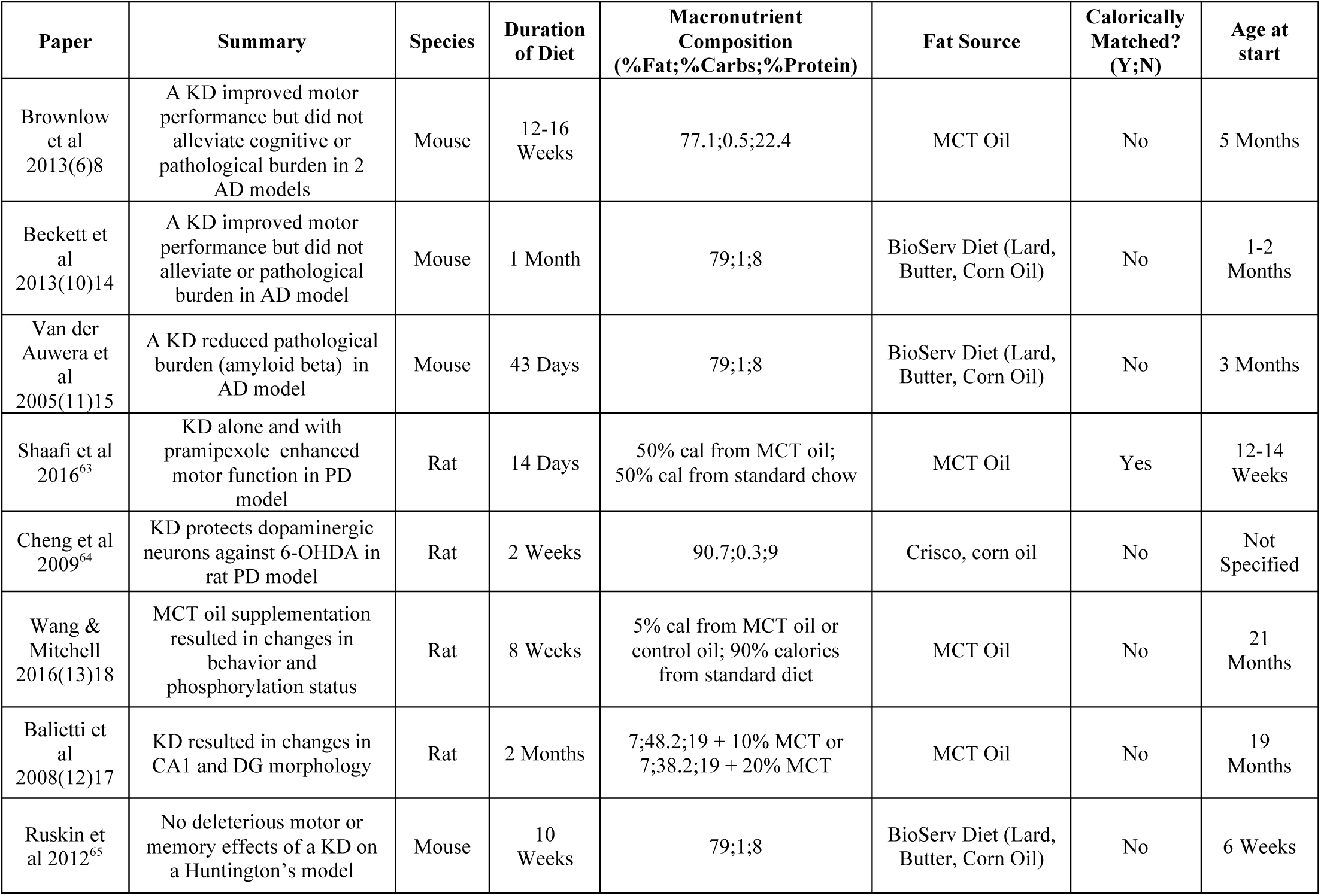
Comparison of variations in previously implemented age- or neurodegenerative disease-related ketogenic diet studies.

While little is known regarding the efficacy of KDs in aged animals, these diet regimens have extensively been shown to be effective at treating seizure disorders(1,2,4). Importantly, epilepsy shares biological features with aging, such as aberrant excitation in the hippocampus (14–17)(17)^17^. Moreover, the incidence of subclinical seizures is higher in individuals with Alzheimer’s disease (18). In fact, anti-epileptic medications have been shown to reduce aberrant hippocampal activity and improve cognitive outcomes in aged rats (14,19) and humans with mild cognitive impairment (20). Because the hippocampus is vulnerable in old age (21,22), shows glucose dysregulation (23,24), and plays a major role in epileptogenesis(25), this structure is likely to exhibit neurobiological alterations following a long-term KD. Thus, as a first pass, it is critical to determine how glucose and monocarboxylate transporter expression within the HPC is altered by a KD and if shifting metabolism to using ketone bodies can reverse reported age-related alterations in the expression of metabolism-related transporters within this brain region (26,27). Furthermore, because neurotransmission is metabolically demanding and the levels of vesicular transporter proteins are known to go down with aging (28,29), it is important to examine whether the KD impacts levels of vesicular transporters in the hippocampus of young and aged animals. The current study aimed to test the impact of a 12-week KD on transporter expression within hippocampus and body composition of young and aged rats.

## Materials and Methods

### Subjects & Handling

Young (4 months; n=26) and aged (20 months: n=30) male Fischer 344 x Brown Norway F1 Hybrid rats from the NIA colony at Taconic Farms were used in this study. Rats were housed individually and maintained on a 12-hr light;dark cycle, and all behavioral testing was performed in the dark phase. Rats were given one week to acclimate to the facility prior to blood measurements. All experimental procedures were performed in accordance with National Institutes of Health guidelines and were approved by Institutional Animal Care and Use Committees at the University of Florida.

### Diet

Prior to diet administration, rats were randomly assigned to either a high fat;low carbohydrate KD (Lab Supply; 5722, Fort Worth, Texas) mixed with MCT oil (Neobee 895, Stephan, Northfield, Illinois) or a calorically and micronutrient comparable control diet (Lab Supply; 1810727, Fort Worth, Texas; see Figure 1 & Supplementary Table 1 for macro and micronutrient composition). There were 13 young and 15 aged rats in each diet group. Because aged animals were used in this study, care was taken to provide adequate protein as well as sufficiently high levels of choline to protect the liver. Rats were weighed daily and given 51 kcal at the same time each day. Access to water was ad libitum and water intake was measured daily in a subset of rats (n = 3;group) to monitor potential effects of the diet on water consumption.

**Figure 1:**
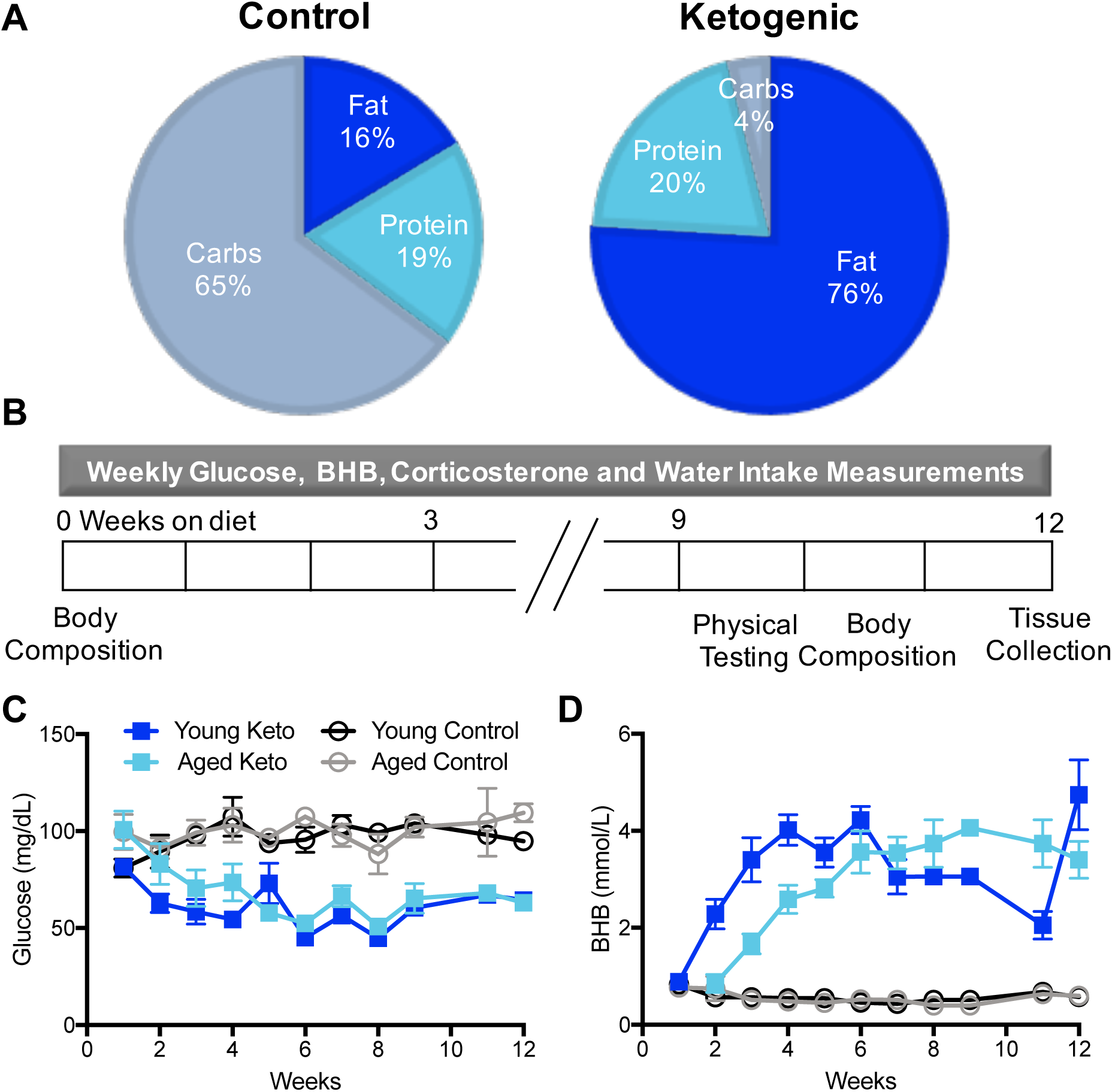
Dietary composition, timeline of experiments, and glucose and ketone body measurements in young and aged rats. (A) Macronutrient composition of the control and ketogenic diets. The ketogenic diet was composed of 76% fat, 20% protein and 4% carbohydrates, whereas the control diet was composed of 16% fat, 19% protein and 65% carbohydrates. Diets were nutrient matched and rats were fed equivalent calories (See Table 1; for micronutrient profile see Supplemental Table 1). (B) Timeline of experiments. Body composition analysis was performed using TD-NMR. Physical testing included body composition analysis as well as rotarod and grip strength tests (Supplemental Fig 1). (C) Diet significantly affected serum glucose levels (p < 0.001), such that they decreased over time in animals on the ketogenic diet regardless of age. (D) Diet significantly affected serum BHB levels (p < 0.001), such that they increased over time in animals on the ketogenic diet regardless of age, although the rate at which young and aged animals reached asymptotic levels of serum BHB significantly differed (p < 0.001), with the aged rats taking longer. Error bars represent ± 1 standard error of the mean (SEM).

### Beta-hydroxybutyrate and glucose measurements

β-hydroxybutyrate (BHB; mmol;L), the prominent ketone body in the blood, and glucose levels (mg;dL) were determined using the Precision Xtra blood monitoring system (Abbott Diabetes Care, Inc, Alameda, CA). Before starting the control or KD, baseline levels were obtained from each rat while free feeding on standard laboratory chow. Rats either underwent blood collection from the saphenous vein (n = 3;group) or received a small tail nick to obtain several drops of blood for monitoring glucose and ketone body levels (n = 10 young, 12 aged rats;diet group). Measurements were taken one hour after eating at multiple weekly time points between weeks 1-12.

### Body composition

Brown and white adipose tissue was isolated (n=8 rats per group) at the time of sacrifice during week 12 by an experimenter blind to diet and age group. Perirenal, retroperitoneal, and epididymal white adipose tissues, as well as interscapular brown adipose tissue (iBAT) were excised and weighed (see Figure 2D). White adipocytes have few mitochondria and are single lipid droplets. In contrast, brown adipocytes consist of many small lipid droplets, are rich of iron and contain mitochondria, causing the identifiable brown discoloration. During the excision of the iBAT, all the muscle and white adipose tissue that were connected to the iBAT were removed, and the remaining tissue was weighed. In 3 rats;group fat was only measured at the time of sacrifice.

**Figure 2:**
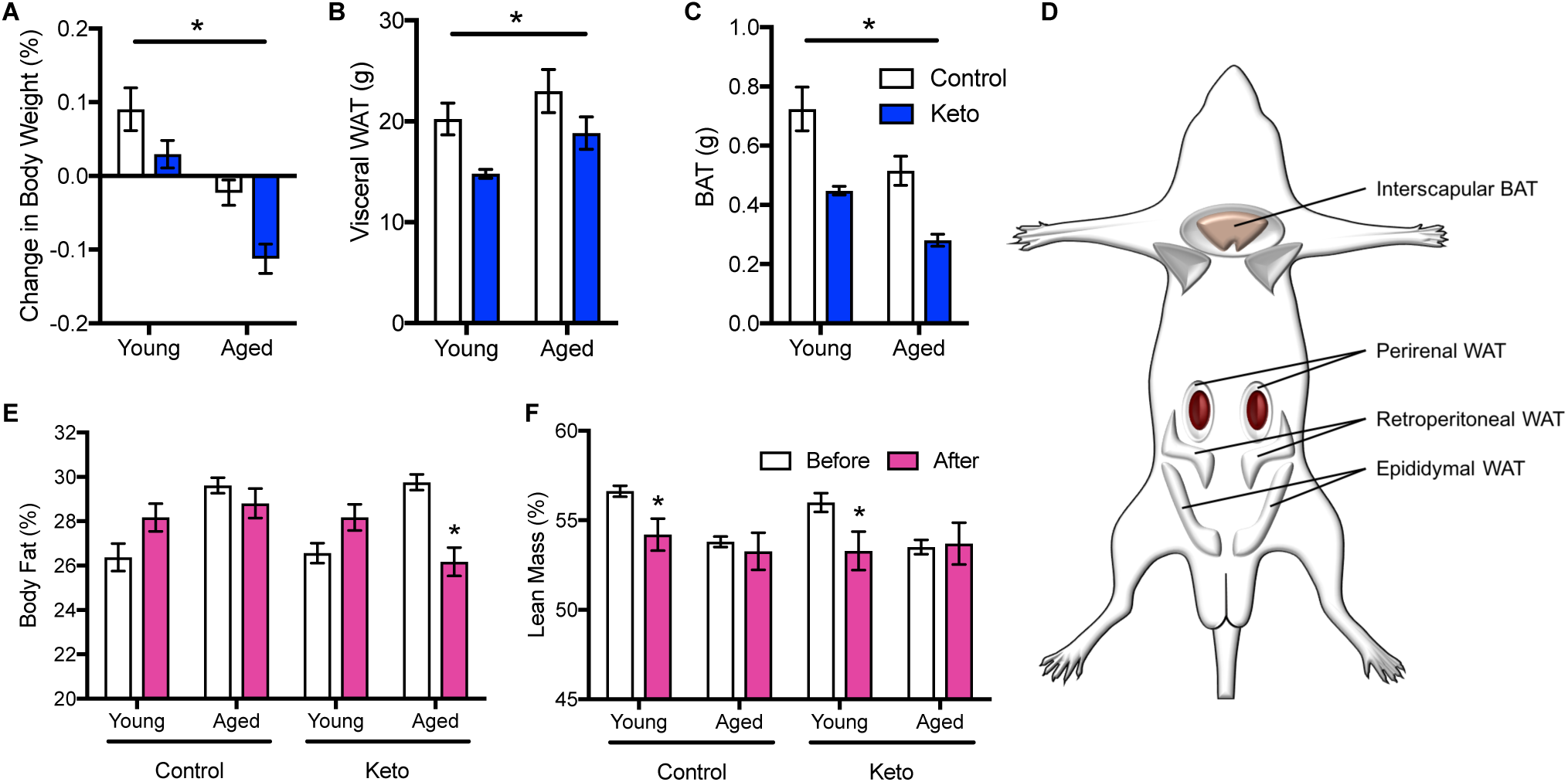
The effects of a ketogenic diet on body composition. (A) There was a significant effect of both age and diet (p <0.001 for both) on the change in body weight after 12 weeks on a ketogenic diet, such that aged rats on a ketogenic diet lost more weight than rats on a calorie-matched control diet. (B) There was a main effect of both age (p < 0.04) and diet (p < 0.01) on the amount of visceral white adipose tissue (WAT) obtained from rats after 12 weeks. (C) Similar to WAT, there was an effect of both age and diet (p < 0.001 for both) on the amount of brown adipose tissue (BAT) recovered. (D) Schematic of brown and white adipose tissue depots quantified. (E) There was a significant main effect of both age (p < 0.001) and diet (p = 0.03) on the change in body fat %, but no significant age by diet interaction (p = 0.07). (F) There was a significant main effect of age (p = 0.01), but not diet (F_[3,40]_ = 0.06; p = 0.81), on the change in lean mass %. Error bars represent ± 1 SEM and asterisks indicate a p value <0.05.

In the other 10 young and 12 aged rats;group time domain-nuclear magnetic resonance (TD-NMR) imaging was performed using a Minispec LF series machine (Bruker; Billerica, Massachusetts) to obtain lean, fat and fluid mass. Measurements were taken before initiation of the diet, and then after 10 weeks on the diet. Rats were weighed, positioned in a clear plastic tube and then placed inside the machine. Each rat was measured twice per session and the average was taken for each rat.

### Tissue Collection

The hippocampus was collected from 5 rats per group. One hour after feeding, rats were anesthetized briefly in a jar of isoflurane and decapitated. Blood was tested for BHB, glucose and corticosterone as described above. The brain was extracted and hemisected, and the left hippocampus was extracted for protein quantification. All tissue samples were immediately frozen on dry ice and stored at −80°C.

### Preparation of hippocampal membrane fractions and Western blotting

Hippocampal tissues was homogenized, and the membrane fractions were isolated according to previously published procedures (30) (see supplemental methods for detail). 5 μg of protein;lane were separated on 4-15% TGX gels (Bio-Rad Laboratories, Hercules, CA, USA) at 200V for 40 minutes in tris-glycine running buffer (Bio-Rad). Total protein was transferred to a 0.45 μm pore nitrocellulose membrane at 20 V for 7 minutes using iBlot Gel Transfer Nitrocellulose Stacks (NR13046-01, Invitrogen, Waltham, MA, USA) and an iBlot machine (Invitrogen, Waltham, MA, USA). All experiments were conducted in triplicate and the loading order of samples was randomized between gels and experiments to control for systematic variation in the electrophoresis and electroblotting procedures.

Immediately after transfer, membranes were stained for total protein using LiCor’s Revert total protein stain for 5 minutes (Li-Cor, 926-11011) and scanned using a 685 nm laser on an Odyssey IR Scanner (Li-Cor, Lincoln, Nebraska USA) to detect total protein levels and then placed into Rockland blocking buffer (Rockland Blocking Buffer, Rockland, Gilbertsville, PA, USA) for 1 hour at room temperature. After blocking membranes were incubated at 4°C overnight with antibodies raised against MCT1, MCT2, MCT4 (ketone body, lactate and pyruvate transporters); GLUT1, GLUT3, GLUT4 (glucose transporters); VGLUT1 (vesicular glutamate transporter 1), or VGAT (vesicular GABA transporter; see Supplemental Table 2 for expanded description of antibodies). Membranes were washed in tris buffered saline before incubation in donkey anti-rabbit or donkey anti-mouse secondary antibodies conjugated to IRDye800 (diluted 1:15000; LICOR). Blots were scanned using a 785 nm laser on an Odyssey IR Scanner. The target protein:total protein ratio was calculated for each technical replicate and data from each independent biological sample were transformed to percent expression of the young, control diet-fed group (i.e. mean of this group is set to 100%) for statistical analysis.

**Table 2:** Macronutrient composition of ketogenic and control diets.

|  | Control Diet | Ketogenic Diet |
| --- | --- | --- |
| **Fat %; kcal** | 16.35%; .7kcal/day | 75.5%; 36.60kcal/day |
| **Protein %;kcal** | 1.76%; .54kcal/day | 20.12%; 10.2kcal/day |
| **Carbohydrates %;kcal** | 64.%; 33.kcal/day | 3.5%; 1.6kcal/day |
| **Total kcal/day** | 51kcal/day | 51kcal/day |

### Statistical Analysis

All quantitative data are represented as mean ± SEM. For all longitudinal data, statistical significance was tested using a repeated measures ANOVA, followed by paired-samples t-tests where appropriate, unless otherwise stated. For endpoint or single time point data, a 2 x 2 multifactorial ANOVA was used, followed by paired- or unpaired-samples t-tests where appropriate, unless otherwise stated. For all factorial ANOVAs each factor had two levels (for diet: control and keto; for age group: young and aged). For all repeated measures ANOVAs the within subjects factor was weeks on diet. The null hypothesis was rejected at the level of p > 0.05 unless it was Bonferroni corrected for multiple comparisons, as stated in the text. All analyses were performed with the Statistical Package for the Social Sciences v24 (IBM, Armonk, NY).

## Results

### The initiation and physiological impact of nutritional ketosis in aged rats

Nutritional ketosis is associated with lower blood glucose and elevated β-hydroxybutyrate (BHB). To demonstrate the efficacy of the KD used here for initiating and maintaining ketosis, glucose and BHB were measured from the blood on a weekly basis (Figure 1). Prior to diet administration, a 2 x 2 factorial ANOVA was used to compare the baseline glucose levels across diet and age groups. There was a main effect of age (F_[1,52]_ = 6.40; p < 0.02), but not diet (F_[1,52]_ = 0.01; p = 0.91), such that all aged animals had higher blood glucose levels than the young group, suggestive of age-associated insulin dysregulation. Repeated-measures ANOVA (within subjects factor of week and between subjects factors of age and diet) of blood glucose levels during weeks 0-11 indicated a significant change over the duration of the diet (F_[9,144]_ = 80.34; p < 0.001; Figure 2C). Moreover, there was a significant main effect of both age (F_[1,16]_ = 28.467; p < 0.001) and diet (F_[1,16]_ = 968.887; p < 0.001) on overall glucose levels. Alterations in glucose levels from baseline through the diet varied significantly as a function of diet group, as indicated by a significant glucose by diet interaction (F_[10,160]_ = 15.488; p < 0.001). Planned contrasts showed that after one week on the diet, there was a significant difference in glucose levels between the KD and control diet groups, and this was maintained for the duration of the diet (p < 0.01 for all comparisons). The interaction effect between blood glucose levels and age was also significant (F_[10,150]_ = 2.675; p < 0.01). Specifically, the young rats on the control diet had glucose levels that increased reaching similar levels to the aged control rats. Conversely, the aged rats on the KD had blood glucose levels that decreased from baseline and became similarly low to young KD rats.

The levels of β-hydroxybutyrate (BHB), the predominant ketone body in the blood, were obtained from the samples collected at baseline and over diet duration (Figure 2D). At baseline, a 2 x 2 factorial ANOVA indicated that BHB did not significantly differ with age (F_[1,52]_ = 0.11; p = 0.74) or diet (F_[1,52]_ = 0.23; p = 0.63). Repeated measures ANOVA for BHB comparing baseline to subsequent weeks indicated a significant change across the 11 weeks (F_[9,144]_ = 4.95; p < 0.001). Moreover, there was a significant main effect of diet on overall BHB (F_[1,16]_ = 467.862; p < 0.001), but not a significant effect of age (F_[1,16]_ = 0.749; p = 0.40), and no significant age by diet interaction (F_[1,16]_ = 1.642; p = 0.218). These data demonstrate that the KD was able to significantly increase BHB levels in both young and aged rats.

Interestingly, changes in BHB levels over the 11 weeks significantly interacted with age group (F_[10,160]_ = 6.69; p < 0.001), suggesting that young and aged rats initiate nutritional ketosis at different rates. This idea was supported by the observation that during the second and third week of the diet, there was a significant age by BHB by diet interaction (p < 0.01 for both comparisons). Specifically, the young rats on the KD had elevated BHB relative to aged rats, but levels in young and aged rats on the control diet did not differ. Importantly, this difference between young and aged rats in the KD group was ameliorated by week 4 (p = 0.651), and remained similar between age groups for the duration of the diet. The extended amount of time required for aged rats to reach asymptotic high levels of blood BHB could be due to the elevated blood glucose and increased body mass they show at baseline, which could be associated with larger glycolysis stores. Finally, there were no changes in corticosterone levels, average water consumption, strength or motor coordination with diet (see supplemental results and Figure S1), indicating that adverse physiological consequences in old animals on the KD were not detected.

Weight was recorded daily for rats and weekly averages were determined. At baseline, a 2 x 2 factorial ANOVA indicated aged rats were significantly heavier than young (F_[1,52]_ = 544.52, p < 0.001), but weight did not significantly differ between diet groups (F_[1,52]_ = 0.10, p = 0.76). Moreover, the interaction between age and diet on weight did not reach statistical significance (F_[1,52]_ = 0.01, p = 0.91). After 12 weeks on the KD, a 2 x 2 factorial ANOVA indicated there was a significant main effect of both age (F_[1,52]_ = 30.81, p < 0.001) and diet (F_[1,52]_ = 10.76, p < 0.01) on change in body weight (Figure 2A), but the age by diet interaction was not significant (F_[1,31]_ = 0.38, p = 0.54). Post hoc analysis on the effect of diet on weight change over 12 weeks within each age group revealed that the aged rats on the KD lost significantly more weight than the aged rats on the control diet (t_[16]_ = 3.40; p < 0.01 corrected *α* = 0.025), while weight change across diet groups in young animals did not significantly differ (t_[15]_ = 1.60; p = 0.13; corrected *α* = 0.025).

The amount of post mortem visceral white adipose tissue (WAT) significantly varied by both age (F_[1,28]_ = 4.74, p < 0.04) and diet (F_[1,28]_ = 9.39, p < 0.01), but the diet by age interaction did not reach statistical significance (F_[1,28]_ = 0.16, p = 0.69; Figure 2B). Additionally, the amount of post mortem brown adipose tissue (BAT) indicated that aged rats had significantly less (F_[1,28]_ = 16.68, p < 0.001). Moreover, BAT was significantly reduced by the KD (F_[1,28]_ = 30.71, p < 0.001), but there was not a significant age by diet interaction (F_[1,28]_ = 0.20, p = 0.66; Figure 2C).

When body composition was measured longitudinally with TD-NMR, there was a significant effect of both age (F_[3,40]_ = 38.49; p < 0.001) and diet (F_[3,40]_ = 4.86; p = 0.03) on the change in body fat %. While the age by diet interaction did not reach statistical significance (F_[3,40]_ = 3.60; p = 0.07; Figure 2E), there was a trend for the aged rats to lose body fat, while the young rats did not show reduced body fat and may actually have had a modest increase over 12 weeks. If fat loss is commensurate with reductions in lean mass, weight loss may not represent improved body conditioning. Importantly, the significant difference on the change in lean mass % (F_[3,40]_ = 7.18; p = 0.01) was due to reductions in young animals while the aged rats maintained lean mass. Moreover, there was no significant effect of diet on change in lean mass % (F_[3,40]_ = 0.06; p = 0.81), nor was there an age by diet interaction (F_[3,40]_ = 0.30; p = 0.58; Figure 2F). These data indicate that the KD does not lead to loss of muscle mass in aged animals and may in fact improve overall body condition.

### Transporter Quantification

All glucose transporters examined had a significant main effect of diet on expression within the hippocampus (p < 0.01 for all; see Supplementary Table 3), such that transporter expression decreased with the KD (Figures 3A & S2). GLUT1, which is found on epithelial cells of the blood brain barrier and some glial cells, was the only glucose transporter in which aged rats had significantly lower expression compared to young (F_[1,15]_ = 7.181; p < 0.02). For GLUT1, there was also a significant interaction of age and diet (F_[1,15]_ = 5.619; p < 0.05). This interaction occurred because the young KD rats had lower levels of GLUT1 compared to young control rats, while levels in aged rats did not differ across diet.

**Figure 3:**
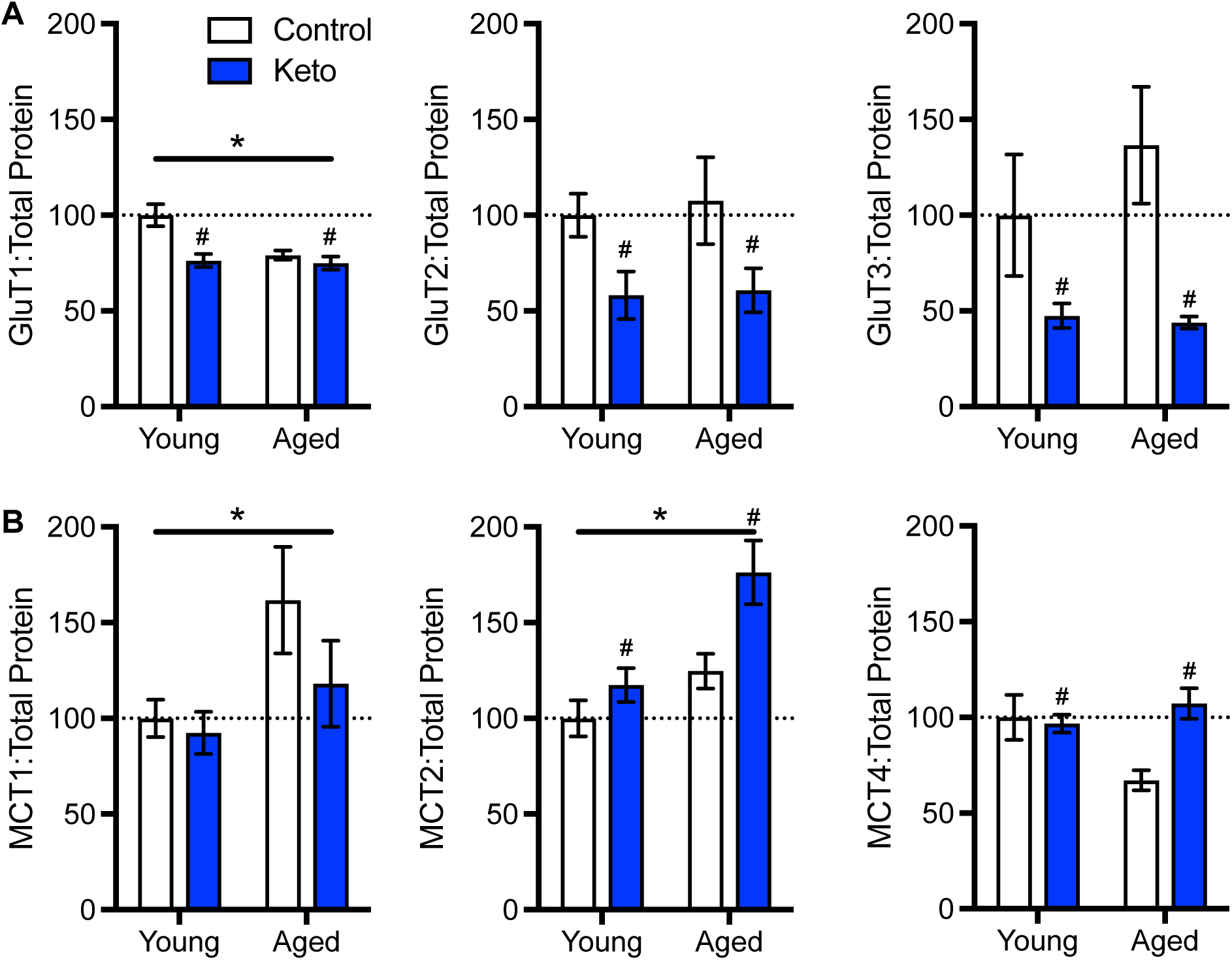
(A) Glucose and (B) monocarboxylate transporter quantification in hippocampal homogenates from young and aged rats on ketogenic and control diets showed age and diet-related alterations in expression level (see Figure S2 for complete gel images). There was a main effect of age on MCT1, MCT2, and GLUT1 (p ≤ 0.04 for all comparisons). There was a main effect of diet on MCT2, MCT4, GLUT1, GLUT2, and GLUT3 (p ≤ 0.05 for all comparisons). There was also a significant age by diet interaction for MCT4 and GLUT1 (p < 0.04 for all comparisons). See Table 3 for complete statistical results. Error bars represent ± 1 SEM, asterisks indicate a p value <0.05 for age and hashtags indicate a p value < 0.05 for diet.

**Table 3:** Effects of age, diet and age by diet interactions on hippocampal monocarboxylate, glucose, vesicular GABA and vesicular glutamate transporter protein expression level in young and aged rats on a control or ketogenic diet.

| Transporter | Age | Diet | Age*Diet |
| --- | --- | --- | --- |
| MCT1 | $F_{[1,15]} = 5.650$ | $F_{[1,15]} = 1.42$ | $F_{[1,15]} = 0.65$ |
| | $p = 0.031$ | $p = 0.184$ | $p = 0.341$ |
| MCT2 | $F_{[1,15]} = 12.446$ | $F_{[1,15]} = 8.473$ | $F_{[1,15]} = 2.075$ |
| | $p = 0.003$ | $p = 0.011$ | $p = 0.170$ |
| MCT4 | $F_{[1,15]} = 1.866$ | $F_{[1,15]} = 5.008$ | $F_{[1,15]} = 6.920$ |
| | $p = 0.192$ | $p = 0.041$ | $p = 0.019$ |
| GLUT1 | $F_{[1,15]} = 7.181$ | $F_{[1,15]} = 11.425$ | $F_{[1,15]} = 5.619$ |
| | $p = 0.017$ | $p = 0.004$ | $p = 0.032$ |
| GLUT2 | $F_{[1,15]} = 0.135$ | $F_{[1,15]} = 0.57$ | $F_{[1,15]} = 0.031$ |
| | $p = 0.728$ | $p = 0.007$ | $p = 0.862$ |
| GLUT3 | $F_{[1,15]} = 0.594$ | $F_{[1,15]} = 11.458$ | $F_{[1,15]} = 0.74$ |
| | $p = 0.453$ | $p = 0.004$ | $p = 0.365$ |
| VGAT | $F_{[1,15]} = 2.23$ | $F_{[1,15]} = 19.802$ | $F_{[1,15]} = 1.184$ |
| | $p = 0.108$ | $p = 0.000$ | $p = 0.294$ |
| VGLUT1 | $F_{[1,15]} = 7.423$ | $F_{[1,15]} = 3.302$ | $F_{[1,15]} = 6.011$ |
| | $p = 0.016$ | $p = 0.089$ | $p = 0.027$ |

MCT1, which is found on epithelial cells on the blood brain barrier as well as some glial cells, had a significant main effect of age (F_[1,15]_ = 5.650; p < 0.05), such that aged rats expressed higher transporter protein levels (Figure 3B). However, there was no main effect of diet (F_[1,15]_ = 1.42; p = 0.18), nor was there an age by diet interaction (F_[1,15]_ = 0.65; p = 0.34). MCT2, which is found on cell bodies and post synaptic terminals of neurons, also had a main effect of age (F_[1,15]_ = 12.446; p < 0.01). Additionally there was a main effect of diet (F_[1,15]_ = 8.473; p = 0.01), such that rats on the KD had decreased transporter levels, but no age by diet interaction. MCT4, located on astrocytes, had a significant main effect of diet (F_[1,15]_ = 5.008; p = 0.04) such that rats on the KD had decreased transporter levels. Although there was no significant age effect (F_[1,15]_ = 1.66; p = 0.19), there was a significant age by diet interaction (F_[1,15]_ = 6.920; p < 0.02). In fact, the KD restored MCT4 expression levels in the aged rats to the levels observed in young controls.

Vesicular transporter expression in the hippocampus for glutamate and GABA were also examined (Figure 4). There was a significant main effect of age on VGLUT1 expression (F_[1,15]_ = 7.423; p < 0.02), but not of diet (F_[1,15]_ = 3.302; p = 0.089). However, the interaction between age and diet was significant (F_[1,15]_ = 6.011; p < 0.03). Critically, this was due to the reversal of age-related declines in VGLUT1 expression present in the control rats by a KD, such that aged rats in nutritional ketosis had levels of hippocampal VGLUT1 that were comparable to young control animals. Lastly, there was a significant main effect of diet for VGAT (F_[1,15]_ = 19.802; p < 0.001), with greater protein levels being observed in the KD rats. There was no age effect (F_[1,15]_ = 2.23; p = 0.11) or age by diet interaction (F_[1,15]_ = 1.12; p = 0.29) for VGAT expression, indicating that animals on a KD expressed higher protein levels than rats on a control diet, irrespective of age.

**Figure 4:**
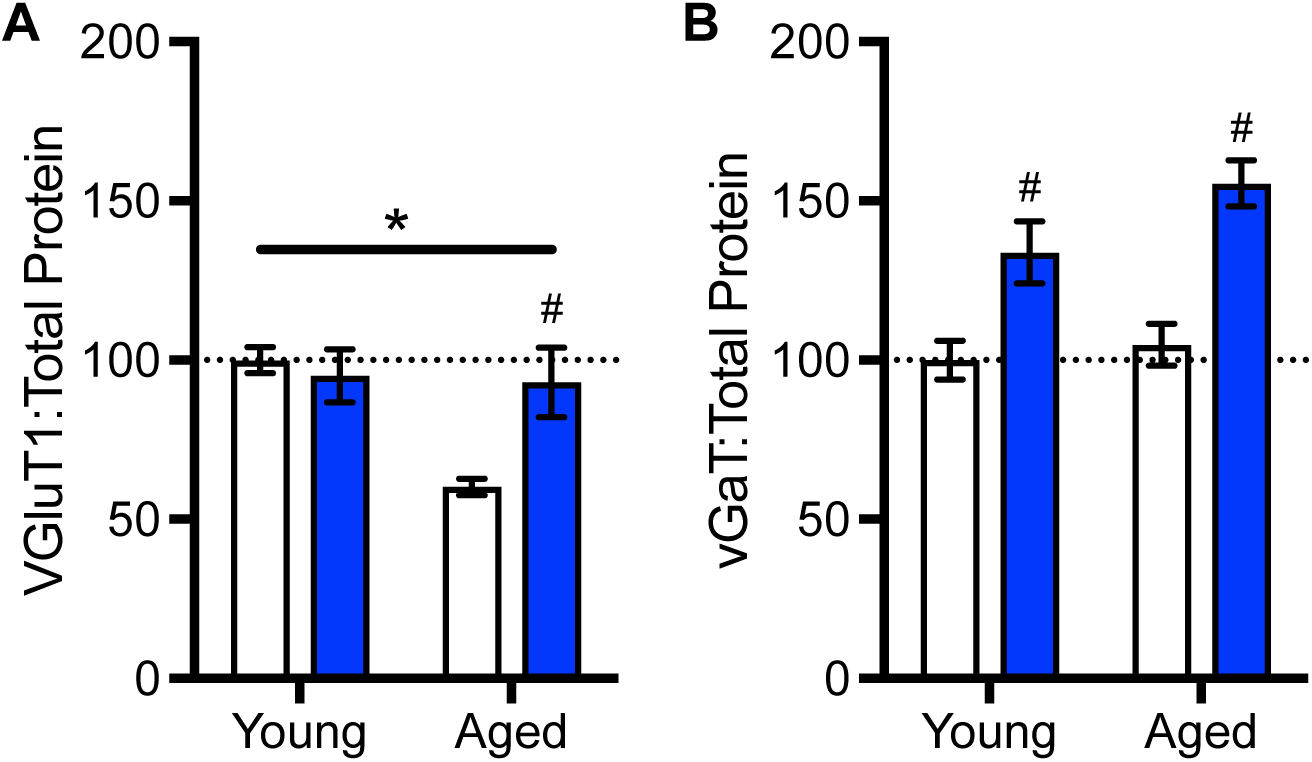
Vesicular (A) glutamate and (B) GABA transporter quantification in hippocampal homogenates from young and aged rats on ketogenic and control diets showed age and diet-related alterations in expression levels (see figure S2 for complete gel images). There was a main effect of age (p ≤ 0.02), and a significant age by diet interaction for VGLUT1 (p < 0.03). There was a significant main effect of diet on VGAT (p ≤ 0.001). See Tables 3 for complete statistical results. Error bars represent ± 1 SEM, asterisks indicate a p value <0.05 for age and hashtags indicate a p value < 0.05 for diet.

## Discussion

Aged individuals are at an increased risk for neurodegenerative diseases, cardiovascular disease, diabetes, cancer, increased oxidative stress and the accumulation of damaged proteins and nucleic acids(8). Recently there has been a call for nutritional cognitive neuroscience research, advocating a dietary intervention for global neurological deficits affecting older adults(31). KDs may be a potential intervention strategy for aged individuals, as glucose metabolism deficits are observed in this population(32,33). Although there are many ongoing studies of the effects of KDs and diets high in oils/fats on age-related disease states, additional data are necessary to establish the safety and efficacy of a KD in older adults.

The current study examined the impact of implementing a long-term KD on several markers of peripheral health and hippocampal biochemistry in a rat model of old age. As varied composition and implementation of KDs complicates the interpretation of existing literature and reconciliation of information regarding health outcomes (See Table 1), the current study took care to ensure that both diets were calorically and nutritionally equivalent, including protein and micronutrient content in young and aged rats. Furthermore, rather than the commonly used long chain fatty acids, such as lard or butter, the current study used medium chain triglyceride (MCT) oil as the fat source, as these are more readily digested and converted into ketone bodies(34).

A critical finding from these data are that when protein and caloric intake are matched between control and KD animals, KD-fed aged rats had decreased glucose and increased ketone body levels, demonstrating the efficacy of the diet at inducing a state of nutritional ketosis at advanced ages. While the levels of ketone bodies in aged rats matched young by week 4, there was significant age by diet interaction for weeks 2 and 3 on the diet. This indicated that aged rats take longer to initiate ketosis. While the mechanism of delayed ketosis in aged rats is unknown, this finding demonstrates that short-term dietary intervention may be insufficient to produce measurable physiological changes in old animals.

One criticism of the KD is the potential for decreased muscle mass and increased muscle fatigue from inadequate protein intake(35), which could be particularly problematic for older individuals at risk for sarcopenia. Importantly, the high BHB levels in all ketogenic rats with normative dietary protein content (∼20%) show that drastic decreases in protein are not necessary for the induction of ketosis. Moreover, the current data show a preservation of strength, coordination and no signs of muscle mass deterioration in aged rats. This finding is consistent with other data showing that ketosis prevents muscle loss in starved individuals(36), and promotes weight loss, with no declines in strength or muscle mass in humans(37). Aged rats on the KD also had larger changes in body weight and composition compared to aged rats on the calorie-matched control diet, which was marked by a decrease in visceral fat and body fat percentage. Importantly, these results were achieved without a reduction in caloric intake relative to controls. Thus, profound calorie restriction is not necessary for weight loss on a KD. These positive changes on body composition are especially important for aged individuals, as they are at increased risk of developing metabolic syndrome, which is linked to cognitive decline (38). In fact high amounts of WAT in aged individuals correlate with lower scores on several types of memory tasks in humans(39), and there is a strong link between unhealthy eating habits and poorer performance on hippocampal dependent tasks in rodents (for review see(40)). Furthermore, it is well established that obesity in mid-to late-life increases the risk for dementia and AD(41).

While reductions in WAT and improved body composition in aged animals are likely to benefit cognition by several possible mechanisms, it is notable that the KD lead to distinct biochemical alterations in the hippocampus of young and aged rats. Consistent with previous data(26), the current study found that aging significantly reduced the levels of GLUT1 expression in the hippocampus, while not affecting levels of GLUT2 and GLUT3. Interestingly, the KD mitigated age-related differences in GLUT1 expression by lowering hippocampal levels of this protein in the young ketogenic animals. The KD also reduced expression of GLUT2 and 3 in the hippocampus of both age groups. While declines in GLUT expression across the brain may be associated with dysfunction in animals on a standard diet, in KD fed rats this likely reflects the lack of reliance on glucose/insulin signaling for neuronal metabolism. Importantly, this is ubiquitous across age groups and matched declines in blood glucose levels.

MCT1 and MCT2 also had significant changes in hippocampal expression levels with age, such that expression of MCT1 was increased and MCT2 was decreased in old control rats. The age-related increase in MCT1 expression may be the result of improper glucose utilization and represent a shift with age towards increased use of monocarboxylates as a fuel source, which has been reported for middle-aged mice and is associated with white matter degeneration(26,27). This difference is ameliorated by a KD, such that expression of MCT1 in young and aged animals on the KD did not differ. The KD increased MCT2 levels in aged rats in a restorative fashion, to be equivalent to levels observed in young control rats. Lastly, there was a significant decrease in MCT4 expression in animals on a KD, regardless of age. Together these data suggest that nutritional ketosis shifts the expression of transporters for different energy substrates in the hippocampus. Conceivably, this could be beneficial to the aged brain, but future cognitive testing in aged rats on a KD will be necessary to test this hypothesis.

Synaptic transmission is particularly demanding of cellular metabolism and vulnerable in advanced age. Therefore, we also measured levels of VGLUT1 and VGAT in the hippocampus. Similar to previous studies(42), there was a significant decrease in VGLUT1 expression with age in control animals. Importantly, the age-associated decline in VGLUT1 was ameliorated by the KD. Because higher levels of VGLUT1 expression are associated with better learning and memory performance(28), this increase suggests that KDs may be able to restore proper neuronal function on old animals.

While there were no changes in VGAT expression with age, this transporter was significantly increased following KD administration in both young and aged rats. Increasing the transport of GABA into vesicles may increase the inhibitory signaling potential of neurons, thus conferring antiepileptic properties unto cells within the hippocampus. Increased quantal release of GABA may serve as one of the mechanisms by which the KD is efficacious in the treatment of refractory epilepsy.

In addition to the positive effects a KD can have on energy metabolism and transporter expression, there are several other mechanisms that may work synergistically to restore normal cognitive function. The moderate reduction in glucose levels in the blood has been suggested to be one possible mechanism of seizure prevention of the KD(43). In fact, a shift towards the utilization of ketone bodies for ATP production has been documented to improve brain functioning during epilepsy, which shares biological features with aging. Moreover, the antiepileptic drug, levetiracetam, has been shown to reduce hyperexcitability in the hippocampus and improve cognitive performance(14,19,20,44). KDs have also been shown to induce the enhancement of energy reserves within the HPC, which can maintain synaptic transmission under periods of metabolic stress(45).

Finally, KDs may be particularly suitable for treating Alzheimer’s disease (AD), as demonstrated by several instances of clinical efficacy in human studies(46). The onset of mild cognitive impairment and Alzheimer’s disease often correlates with epileptiform activity(18), which has shown to be remediated in many studies of epilepsy patients on a KD(1,2). Furthermore, 80% of patients with AD also present with symptoms of diabetes type II or impaired fasting glucose levels(47), which are also positively affected by KDs. The connection with impaired glucose signaling in AD has led to the suggestion that the disease could be reclassified as type III diabetes(48). Furthermore, studies of AD brains have proven that while glucose metabolism is impaired in these patients, ketone body metabolism is not(49,50). Prior studies using young rodent models have shown the feasibility of doing this type of research in animal studies, and here we demonstrate the translational potential of a KD to improve somatic and brain health in an aged population.

---

**Supplementary Figure 1:**
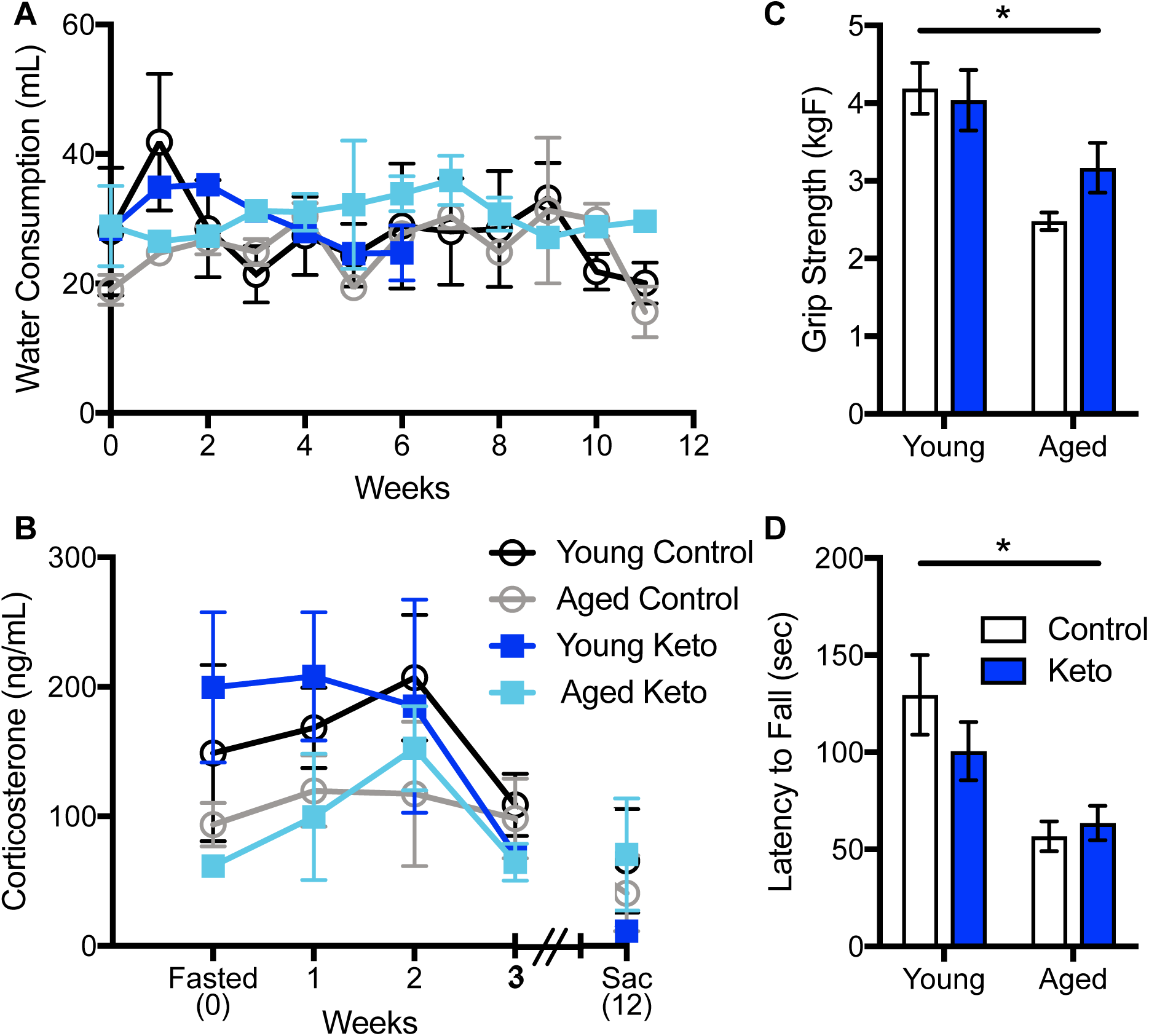
There were no detected adverse side effects of the ketogenic diet in aged rats. (A) There was not a significant effect of age (p = 0.12) or diet (p = 0.65) on corticosterone levels, but levels decreased in all groups throughout the duration of the experiment (p < 0.04). (B) There were no significant effects of diet or age on water consumption (p > 0.2 for both comparisons). (C) There was a significant main effect of age (p < 0.001) but not diet (p = 0.41) on the maximum latency to fall during a rotarod task following 10 weeks of the KD or control diet. (D) There was a significant main effect of age (p < 0.001) but not diet (p = 0.34) on the maximum grip strength after 10 weeks on the KD or control diet. Error bars represent ± standard error of the mean (SEM).

**Supplementary Figure 2:**
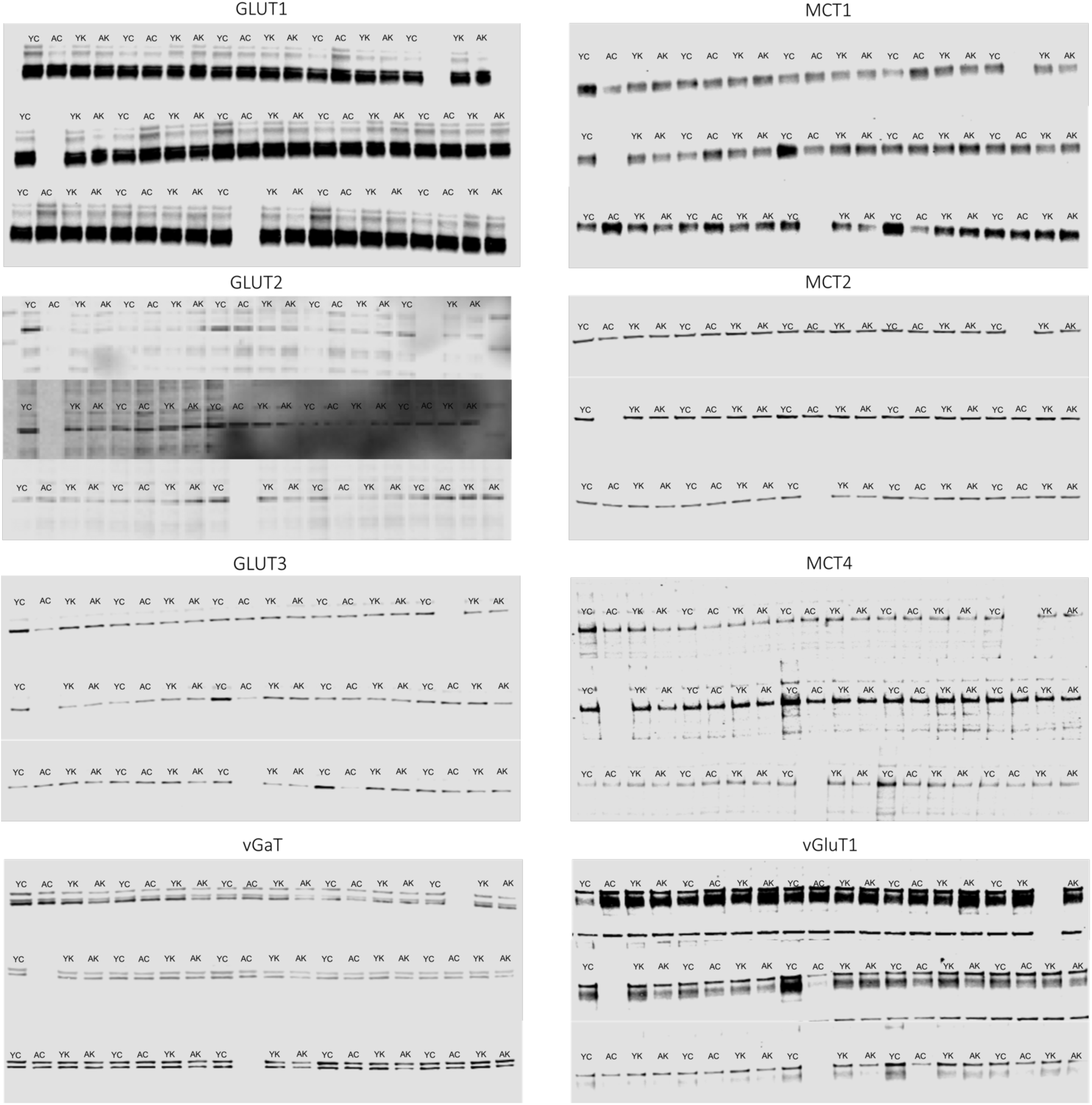
Glucose, monocarboxylate, and vesicular neurotransmitter transporter western blots, run in triplicate. YC = Young control; AC = Aged Control; YK = Young Keto; AK = Aged Keto.

**Supplementary Table 1:** Micronutrient composition of ketogenic and control diets. Diets were supplemented with all essential vitamins and nutrients necessary for proper rodent health and additional choline to support liver health.

|  | **Control Diet** | **Ketogenic Diet** |
| --- | --- | --- |
| **Minerals** |  |  |
| Calcium, % | 0.78 | 0.78 |
| Phosphorus, % | 0.65 | 0.65 |
| Potassium, % | 0.77 | 0.77 |
| Magnesium, % | 0.07 | 0.07 |
| Sodium, % | 0.14 | 0.14 |
| Chloride, % | 0.23 | 0.23 |
| Fluorine, ppm | 1.2 | 1.2 |
| Iron, ppm | 65 | 65 |
| Zinc, ppm | 48 | 48 |
| Manganese, ppm | 76 | 76 |
| Copper, ppm | 7.8 | 7.8 |
| Iodine, ppm | 0.27 | 0.27 |
| Chromium, ppm | 2.6 | 2.6 |
| Molybdenum, ppm | 2.11 | 2.11 |
| Selenium, ppm | 0.32 | 0.32 |
| **Vitamins** |  |  |
| Vitamin A, IU/g | 5.2 | 5.2 |
| Vitamin D-3, IU/g | 1.3 | 1.3 |
| Vitamin E, IU/kg | 68.3 | 68.3 |
| Vitamin K, ppm | 0.65 | 0.65 |
| Thiamin HCl, ppm | 7.9 | 7.9 |
| Riboflavinm, ppm | 8.9 | 8.9 |
| Niacin, ppm | 39 | 39 |
| Pantothenic Acid, ppm | 21 | 21 |
| Folic Acid, ppm | 2.8 | 2.8 |
| Pyridoxine, ppm | 7.5 | 7.5 |
| Biotin, ppm | 0.3 | 0.3 |
| Vitamin B-12, mcg/kg | 19 | 19 |
| Choline, ppm | 1,290 | 1,290 |

**Supplementary Table 2:**
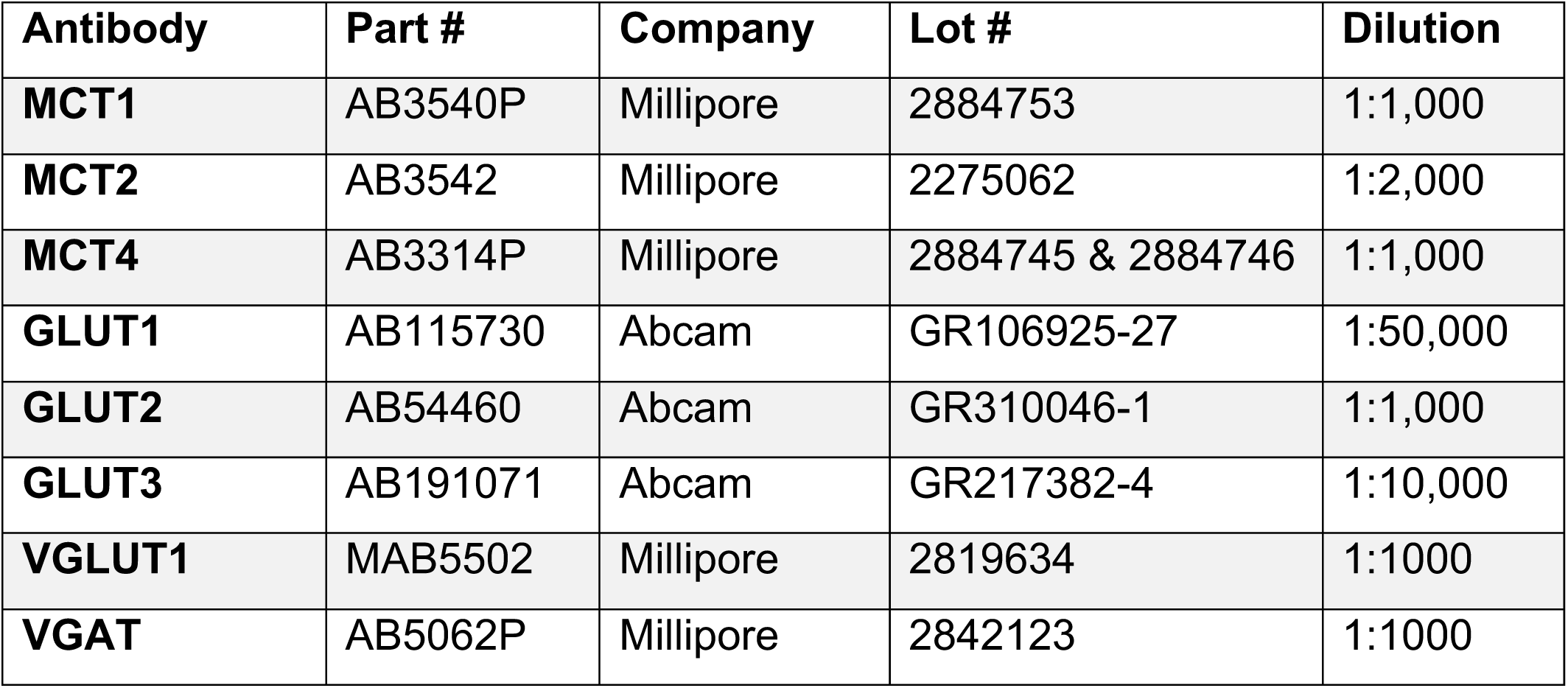
Antibodies used for transporter protein quantification.

